# FAVITES: simultaneous simulation of transmission networks, phylogenetic trees, and sequences

**DOI:** 10.1101/297267

**Authors:** Niema Moshiri, Manon Ragonnet-Cronin, Joel O. Wertheim, Siavash Mirarab

**Affiliations:** Bioinformatics and Systems Biology Graduate Program, UC San Diego, La Jolla, 92093, USA; Department of Medicine, UC San Diego, La Jolla, 92093, USA; Department of Electrical and Computer Engineering, UC San Diego, La Jolla, 92093, USA

## Abstract

**Motivation:** The ability to simulate epidemics as a function of model parameters allows insights that are unobtainable from real datasets. Further, reconstructing transmission networks for fast-evolving viruses like HIV may have the potential to greatly enhance epidemic intervention, but transmission network reconstruction methods have been inadequately studied, largely because it is difficult to obtain “truth” sets on which to test them and properly measure their performance.

**Results:** We introduce FAVITES, a robust framework for simulating realistic datasets for epidemics that are caused by fast-evolving pathogens like HIV. FAVITES creates a generative model to produce contact networks, transmission networks, phylogenetic trees, and sequence datasets, and to add error to the data. FAVITES is designed to be extensible by dividing the generative model into modules, each of which is expressed as a fixed API that can be implemented using various models. We use FAVITES to simulate HIV datasets and study the realism of the simulated datasets. We then use the simulated data to study the impact of the increased treatment efforts on epidemiological outcomes. We also study two transmission network reconstruction methods and their effectiveness in detecting fast-growing clusters.

**Availability and implementation:** FAVITES is available at https://github.com/niemasd/FAVITES, and a Docker image can be found on DockerHub (https://hub.docker.com/r/niemasd/favites).

## 1 Introduction

The spread of many infectious diseases is driven by social and sexual networks (Kelly *et al.*, 1991), and reconstructing their transmission histories from molecular data may be able to enhance intervention. For example, network-based statistics for measuring the effects of Antiretroviral Therapy (ART) in Human Immunodeficiency Virus (HIV) can yield increased statistical power (Wertheim *et al.*, 2011); the analysis of the growth of HIV infection clusters can yield actionable epidemiological information for disease control (Shargie and Lindtjørn, 2007); transmission-aware models can be used to infer HIV evolutionary rates (Vrancken *et al.*, 2014).

A series of events in which an infected individual infects another individual can be shown as a *transmission network*, which itself is a subset of a *contact network*, a graph in which nodes represent individuals and edges represent contacts (e.g. sexual) between pairs of individuals. If the pathogens of the infected individuals are sequenced, which is the standard of HIV care in many developed countries, one can attempt to reconstruct the transmission network (or its main features) using molecular data. Some viruses, such as HIV, evolve quickly, and the phylogenetic relationships between viruses are reflective of transmission histories (Leitner *et al.*, 1996), albeit imperfectly (Ypma *et al.*, 2013; Romero-Severson *et al.*, 2014; Leitner and Romero-Severson, 2018).

Recently, multiple methods have been developed to infer properties of transmission networks from molecular data (Prosperi *et al.*, 2011; Ragonnet-Cronin *et al.*, 2013; Kosakovsky Pond *et al.*, 2018). Efforts have been made to characterize and understand the promise and limitations of these methods: it is suggested that, when combined with clinical and epidemiological data, these methods can provide critical information about drug resistance, associations between sociodemographic characteristics, viral spread within populations, and the time scales over which viral epidemics occur (Grabowski and Redd, 2014). More recently, these methods have become widely used at both local (Campbell *et al.*, 2017) and global scale (Wertheim *et al.*, 2014). Nevertheless, several questions remain to be fully answered regarding the performance of these methods. It is not always clear which method/setting combination performs best for a specific downstream use-case or for specific epidemiological conditions. More broadly, the effectiveness of these methods in helping achieve public health goals is the subject of ongoing clinical and theoretical research.

Accuracy of transmission networks is difficult to assess because the true order of transmissions is not known. Moreover, predicting the impact of parameters of interest (e.g., rate of treatment) on the epidemiological outcomes is difficult. In simulations, in contrast, the ground truth is known and parameters can be easily controlled. The simulation of transmission networks needs to combine models of social network, transmission, evolution, and ideally sampling biases and errors (Villandre *et al.*, 2016).

We introduce FAVITES (FrAmework for VIral Transmission and Evolution Simulation), which can simulate numerous models of contact networks, viral transmission, phylogenetic and sequence evolution, data (sub)sampling, and real-world data perturbations, and which was built to be flexible such that users can seamlessly plug in statistical models at every step of the simulation process. Previous attempts to create an epidemic simulation tool include `epinet` (Groendyke *et al.*, 2012), TreeSim (Stadler and Bonhoeffer, 2013), `outbreaker` (Jombart *et al.*, 2014), `seedy` (Worby and Read, 2015), and `PANGEA.HIV.sim` (Ratmann *et al.*, 2017). A detailed comparison of FAVITES with these tools can be found in Table S1. One key distinction is that FAVITES simulates the full end-to-end epidemic dataset (social contact network, transmission history, incomplete sampling, viral phylogeny, error-free sequences, and real-world sequencing imperfections), whereas each existing tool simulates only a subset of these steps. Another key distinction is that FAVITES allows the user to choose among several models at each step of the simulation, whereas the existing tools are restricted to specific models. After describing the FAVITES framework, we compare its output to real data on a series of experiments, study the properties of HIV epidemics as functions of various model and parameter choices, and finally perform simulation experiments to study two transmission network reconstruction methods.

## 2 Materials and methods

### 2.1 FAVITES simulation process

FAVITES provides a workflow for the simulation of viral transmission networks, phylogenetic trees, and sequence data (Fig. 1). It breaks the simulation process into a series of interactions between abstract modules, and users can select the module implementations appropriate to their specific context. In the statistical sense, the end-to-end process creates a complex composite generative model, each module is a template for a sub-model of a larger model, and different implementations of each module correspond to different statistical sub-models. Thus, the FAVITES workflow does not explicitly make model choices: each module *implementation* makes those choices. The model for a FAVITES execution is defined by the set of module implementations chosen by the user.

**Figure 1:**
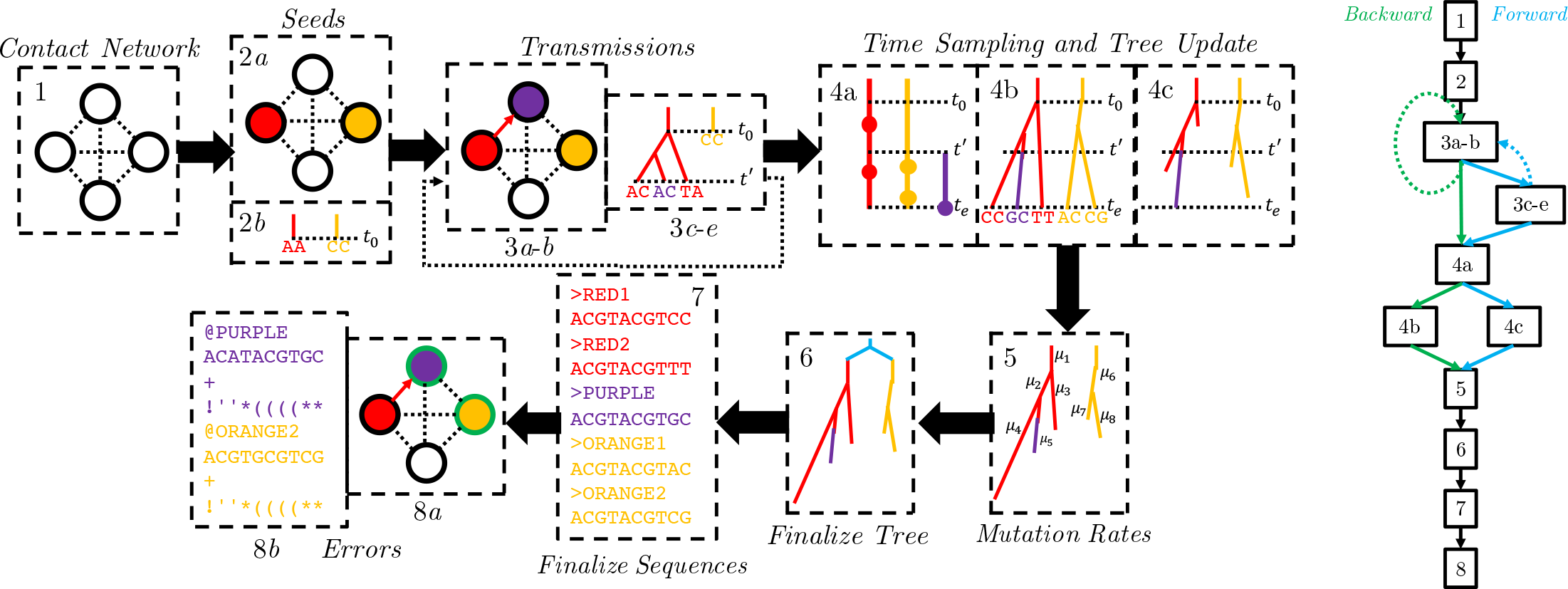
FAVITES workflow. (1) The contact network is generated (nodes: individuals; edges: contacts). (2) *Seed* individuals who are infected at time 0 are selected (2*a*), and a viral sequence is chosen for each (2*b*). (3) The epidemic yields a series of transmission events in which the time of the next transmission is chosen (3*a*), the source and target individuals are chosen (3*b*), the viral phylogeny in the source node is evolved to the transmission time (3*c*), viral sequences in the source node are evolved to the transmission time (*3d*), and a viral lineage is chosen to be transmitted from source to destination (3*e*). Step (3) repeats until the end criterion is met. Step 3*c*-3*e* are optional, as tree and sequence generation can be delayed to later steps. (4) Infected individuals are sampled such that viral sequencing times are chosen for each infected individual (4*a*), viral phylogenies (one per seed) are evolved to the end time of the simulation (4*b*), and viral phylogenies (one per seed) are pruned to reflect the viral sequencing times selected (4*c*). (5) Mutation rates are introduced along the branches of the viral phylogenies and the tree is scaled to the unit of expected mutations. (6) The seed trees are merged using a seed tree (cyan). (7) Viral sequences obtained from each infected individual are finalized. (8) Real-world errors are introduced on the error-free data, such as subsampling of the sequenced individuals (marked as green) (8*a*) and the introduction of sequencing errors (8*b*). The workflows of a typical forward (blue) and backward (green) simulation are shown as well.

FAVITES defines APIs for each module and lets implementation decide how to achieve the goal of the module. The APIs allow various forms of interaction between modules, which enable sub-models that are described as conditional distributions (via dependence on preceding steps) or as joint distributions (via joint implementation). Module implementations can simply wrap around existing tools, allowing for significant code reuse. The available implementations for each step are continuously updated; the full documentation of these implementations can be found online, and a list of current implementations can also be found in the supplemental file.

Simulations start at time zero and continue until a user-specified stopping criterion is met. Error-free and error-prone transmission networks, phylogenetic trees, and sequences are output at the end. FAVITES has eight steps (Fig. 1) detailed below. Depending on the specific implementations, some of the steps may not be needed (we mark these with an asterisk), especially when the phylogeny is simulated backward in time. Also note that steps and modules are not the same; a module may be used in several steps and a step may require multiple modules.

#### Step 1: Contact Network

The *ContactNetworkGenerator* module generates a contact network; vertices represent individuals, and edges represent contacts between them that can lead to disease transmission (e.g. sexual). Graphs can be created stochastically using existing models (Karoński, 1982), including those that capture properties of real social networks (Watts and Strogatz, 1998; Barabási and Albert, 1999; Newman *et al.*, 2002) and those that include communities (Watts, 1999; Fortunato, 2010). For example, the Erdős-Rényi (ER) model (Bollobas, 1984) generates graphs with randomly-placed edges, the Random Partition model (Fortunato, 2010) generates communities, the Barabási-Albert model (Barabási and Albert, 1999) generates scale-free networks whose degree distributions follow power-law (suitable for social and sexual contact networks), the Caveman model (Watts, 1999) and its variations (Fortunato, 2010) generate small-world networks, the Watts-Strogatz model (Watts and Strogatz, 1998) generates small-world networks with short average path lengths, and Complete graphs connect all pairs of individuals (suitable for some communicable diseases). We currently have many models implemented by wrapping around the NetworkX package (Hagberg *et al.*, 2008). In addition, a user-specified network can be used.

#### Step 2: Seeds

The transmission network is initialized in two steps. *a*) The *SeedSelection* module chooses the “seed” nodes: individuals who are infected at time zero of the simulation. *b*\*) For each selected seed node, the *SeedSequence* module can generate an initial viral sequence.

Seed selection has many implemented models, including uniform random selection, degree-weighted random selection, and models that place seeds in close proximity. Seed sequences can be user-specified or randomly sampled from probabilistic distributions. To enable seed sequences that emulate the virus of interest, we implement a model that uses HMMER (Eddy, 1998) to sample each seed sequence from a profile Hidden Markov Model (HMM) specific to the virus of interest. Profile HMMs are appropriate for sampling random sequences that are intended to resemble real sequences because they define a probabilistic distribution over the space of sequences, they can be flexible to insertions and deletions, and they can be sampled in a computationally efficient manner. We provide a set of such prebuilt profile HMMs constructed from multiple sequence alignments (MSAs) of viral sequences.

When multiple seeds are chosen, we need to model their phylogenetic relationship as well. Thus, we also have a model that samples a *single* sequence from a viral profile HMM using HMMER, simulates a *seed tree* with a single leaf per seed individual (e.g. using Kingman coalescent or birth-death models using DendroPy (Sukumaran and Holder, 2010)), and then evolves the viral sequence down the tree to generate seed sequences using Seq-Gen (Rambaut and Grass, 1997).

#### Step 3: Transmissions

An iterative series of transmission events occurs under a transmission model until the *EndCriteria* module triggers termination (e.g. after a user-specified time or a user-specified number of transmission events). Each transmission event has five components.

*a*) The *TransmissionTimeSample* module chooses the time at which the next transmission event will occur and advances the *current* time accordingly, and b) the *TransmissionNodeSample* module chooses a source node and target node to be involved in the next transmission event. These two modules are often jointly implemented. Some of the current implementations use simple models such as drawing transmission times from an exponential distribution and selecting nodes uniformly at random. Others are more realistic and use Markov processes in which individuals start in some state (e.g. Susceptible) and transition between states of the model (e.g. Infected) over time. These Markov models are defined by two sets of transition rates: *nodal* and *edge-based*. Nodal transition rates are rates that are independent of interactions with neighbors (e.g. the transition rate from Infected to Recovered), whereas edge-based transition rates are the rate of transitioning from one state to another given that a single neighbor is in a given state (e.g. the transition rate from Susceptible to Infected given that a neighbor is Infected). The rate at which a specific node *u* transitions from state *a* to state *b* is the nodal transition rate from *a* to *b* plus the sum of the edge-based transition rate from *a* to *b* given neighbor *υ*’s state for all neighbors *υ*. We use GEMF (Sahneh *et al.*, 2017) to implement many compartmental epidemiological models in this manner, including sophisticated HIV models like the Granich *et al.* (2009) model and the HPTN 071 (PopART) model (Cori *et al.*, 2014).
*c*\*) For forward models of tree evolution, the *NodeEvolution* module evolves viral phylogenetic trees of the source node to the current time using stochastic models of tree evolution (Hartmann *et al.*, 2010). We use DendroPy (Sukumaran and Holder, 2010) for birth-death and use our own implementation of dual-birth (Moshiri and Mirarab, 2018) and Yule. Backward-in-time models of evolution (e.g. coalescent), can skip *step 3c*.
*d*\*) The *SequenceEvolution* module is invoked to evolve all viral sequences in the source node to the current time. Commonly-used models of DNA evolution including General Time-Reversible (GTR) model (Tavaré, 1986), and its reductions such as Jukes and Cantor (1969), Kimura (1980), Felsenstein (1981), and Tamura and Nei (1993), are currently available as implementations of *SequenceEvolution*. FAVITES also includes the GTR+Γ model, which incorporates rates-across-sites variation (Yang, 1994). It also includes multiple codon-aware extensions of the GTR model, such as mechanistic (Zaheri *et al.*, 2014) and empirical (Kosiol *et al.*, 2007) codon models. These modules internally use Seq-Gen (Rambaut and Grass, 1997) and Pyvolve (Spielman and Wilke, 2015). When sequences are generated in this step, because viral phylogenies are in units of time, a per-time mutation rate must be assumed by the *SequenceEvolution* module.
*e*\*) The *SourceSample* module chooses the viral lineage(s) in the source node to be transmitted.

Substeps *c* − *e* are required only if the choice of transmission events after time *t* depends on the past phylogeny or sequences up to time *t*. If the choice of future transmission recipients/donors and transmission times are agnostic to past phylogenies and sequences, these tasks can be delayed until later in the workflow, namely in *Steps 4b* and *7*.

#### Step 4: Time Sampling and Tree Update

The patient sampling (i.e., sequencing) events are determined and phylogenetic trees are updated accordingly. Three sub-steps are involved.

*a*) For each individual, the *NumTimeSample* module chooses the number of sequencing times (e.g. a fixed number or a number sampled from a Poisson distribution), the *TimeSample* module chooses the corresponding sequencing time(s) (e.g. by draws from uniform or truncated Gaussian or Exponential distributions, or by sampling right before the first transition of a person to a treated state), and the *NumBranchSample* module chooses how many viral lineages will be sampled at each sequencing time (e.g., single). A given individual may not be sampled at all, thus simulating incomplete epidemiological sampling efforts.
*b*\*) The *NodeEvolution* module is called to simulate the phylogenetic trees *given sampling times*. This step can be used *instead of Step 3c* to evolve only lineages that are sampled, thereby reducing computational overhead. In particular, if the tree simulation model is backwards (e.g. coalescent models), *Step 3c* should be ignored, and the full backward simulation process can be performed at once here. We use VirusTreeSimulator (Ratmann *et al.*, 2017) for coalescent models with constant, exponentially-growing, or logistically-growing population size.
*c*\*) If the tree is simulated in *Step 3c*, it may need to be pruned to only include lineages that are sampled, which we perform here.

#### Step 5: Mutation Rates

Trees from *Step 4* have branch lengths in units of time. To generate sequences, rates of evolution must be assumed. The *TreeUnit* module determines such rates. For example it may use constant rates or may draw from a distribution (e.g., Gamma). Applying rates on the tree yields a tree with branch lengths in units of per-site expected number of mutations. Note that sequences could have been simulated forward in time in *Step 3d*, at which point phylogenies were in units of time. In this case, a joint implementation of the *TreeUnit* and *SequenceEvolution* modules must be used such that per-time mutation rates are chosen in *Step 3d*, and the same mutation rates are used to scale the tree here.

#### Step 6*: Finalize Tree

We now have a single tree per seed. Some implementations of *SeedSequence* also simulate a tree connecting seeds; so the roots of per-seed trees have a phylogenetic relationship. In this case, this step merges all phylogenetic trees into a single global tree by placing each individual tree’s root at its corresponding leaf in the seed tree (Fig. 1).

#### Step 7: Finalize Sequences

The *SequenceEvolution* module is called again. Two scenarios are possible. If sequences have been simulated in *Step 3d*, this step is used to evolve all sequences between the last transmission time and the sampling times. Alternatively, *Step 3d* may have been skipped (as discussed before) and sequence evolution may have been delayed until this point. In that scenario, the *SequenceEvolution* module performs the full sequence simulation on the final tree(s) at once in this step.

#### Step 8: Errors

Error-free data are now at hand. Noise is introduced onto the complete error-free data in two ways.

*a*\*) The *NodeAvailability* module further subsamples the individuals to simulate lack of accessibility to certain datasets. Note that whether or not an individual is sampled is a function of two different modules: *NodeAvailability* and *NumTimeSample* (if *NumTimeSample* returned 0, the individual is not sampled). Conceptually, *NumTimeSample* can be used to model when people are sequenced, while *NodeAvailability* can be used to model patterns of data availability (e.g. sharing of data between clinics).
*b*) The *Sequencing* module simulates sequencing error on the simulated sequences. In addition to sequencing machine errors, this can incorporate other real-world sequencing issues, e.g. taking the consensus sequence of a sample and introducing of ambiguous characters. FAVITES currently uses existing tools to simulate Illumina, Roche 454, SOLiD, Ion Torrent, and Sanger sequencing (Huang *et al.*, 2012; Angly *et al.*, 2012), including support for ambiguous characters.

#### Model validation

We provide tools to validate FAVITES outputs, by comparing the simulation results against real data the user may have (e.g., networks, phylogenetic trees, or sequence data) using various summary statistics (Table S2). In addition to validation scripts, we have several helper scripts to implement tasks that are likely common to downstream use of FAVITES output (Table S3).

### 2.2 Experimental setup

We have performed a set of simulations using the FAVITES framework. In these studies, we compare the simulated data against real HIV datasets, study properties of the epidemic as a function of the parameters of the underlying generative models, and compare two transmission cluster inference tools when applied to sequence data generated by FAVITES. All datasets can be found at https://gitlab.com/niemasd/favites-paper-final.

#### 2.2.1 The simulation model

We selected a set of “base” simulation models and parameters and also performed experiments in which they were varied. For each parameter set, we ran 10 simulation replicates. The base simulation parameters were chosen to emulate HIV transmission in San Diego from 2005 to 2014 to the extent possible. In addition, to show the applicability of FAVITES to other settings, we also performed a simulation with parameters learned from the HIV epidemic in Uganda from 2005 to 2014. For both datasets, we estimate some parameters from real datasets while we rely on the literature where such data are not available. We first describe base parameters for San Diego and then present changing parameters and Uganda parameters (see Tables S4 and S5 for the full list of parameters).

##### Contact network

The contact network includes 100,000 individuals to approximate the at-risk community of San Diego. We set the base expected degree 𝔼_*d*_) to 4 edges (i.e., sexual partners over 10 years). This number is motivated by estimates from the literature (e.g. ≈3 in Wertheim *et al.* (2017) and 3-4 in Rosenberg *et al.* (2011)), and it is varied in the experiments. We chose the Barabási-Albert (BA) model as the base network model because it can generate power-law degree distributions (Barabási and Albert, 1999), a property commonly assumed of sexual networks (Hamilton *et al.*, 2008).

##### Seeds

We chose 15,000 total infected seed individuals uniformly at random based on the estimate of total HIV cases in San Diego as of 2004 (Macchione *et al.*, 2015a).

##### Epidemiological model

We model HIV transmission as a Markov chain epidemic model (see Section 2.1) with states Susceptible (S), Acute Untreated (AU), Acute Treated (AT), Chronic Untreated (CU), and Chronic Treated (CT). All seed individuals start in AU, and transmissions occur with rates that depend for each individual on the number of neighbors it has in each state (Fig. 2). Note that this model is a simplification of the model used by Granich *et al.* (2009).

**Figure 2:**
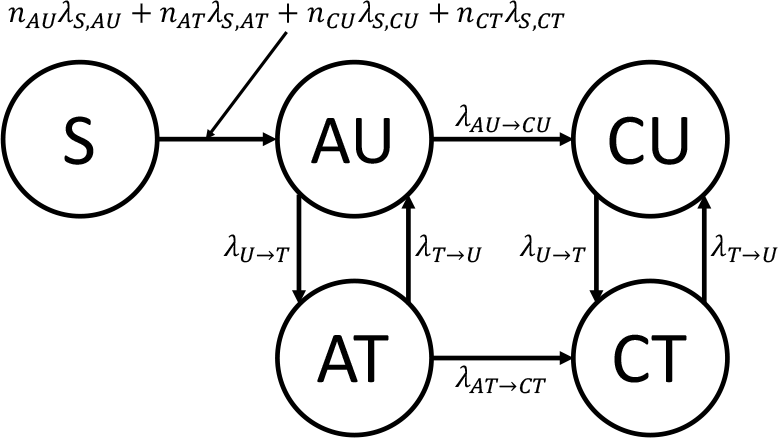
Epidemiological model of HIV transmission with states Susceptible (S), Acute Untreated (AU), Acute Treated (AT), Chronic Untreated (CU), and Chronic Treated (CT). The model is parameterized by the rates of infectiousness of AU (λ_*S*,*AU*_), AT (λ_*S*,*AT*_), CU (λ_*S*,*CU*_), CT (λ_*S*,*CT*_) individuals, and by the rate to transition from AU to CU (λ_*AU*→*CU*_), the rate to transition from AT to CT (λ_*AT*→*CT*_), the rate to start ART (λ_*U*→*T*_), and the rate to stop ART (λ_*T*→*U*_)

We set λ_*AU*→*CU*_ such that the expected time to transition from AU to CU is 6 weeks (Bellan *et al.*, 2015) and set λ_*AT*→*CT*_ such that the expected time to transition from AT to CT is 12 weeks (Cohen *et al.*, 2011). We set λ_*U*→*T*_ such that the expected time to start ART is 1 year from initial infection (O’Brien and Markowitz, 2012), and we define 𝔼_*ART*_= 1/λ_*U*→*T*_ We set λ_*U*→*T*_ such that the expected time to stop ART is 25 months from initial treatment (Nosyk *et al.*, 2015). For the rates of infection λ_*S*,*J*_ for *j* ∈ {*AU*, *CU*, *AT*, *CT*}, using the infectiousness of CU individuals as a baseline, we set the parameters such that AU individuals are 5 times as infectious (Wawer *et al.*, 2005) and CT individuals are not infectious (i.e., rate of 0). Cohen *et al.* (2011) found a 0.04 hazard ratio when comparing linked HIV transmissions between an early-therapy group and a late-therapy group, so we estimated AT individuals to be 1/20 the infectiousness of CU individuals. We then scaled these relative rates so that the total number of new cases over the span of the 10 years was roughly 6,000 (Macchione *et al.*, 2015b), yielding λ_*S*,*AU*_ = 0.1125.

##### Phylogeny

We estimate parameters related to phylogeny and sequences from real data. We used a multiple sequence alignment (MSA) of 674 HIV-1 subtype B *pol* sequences from San Diego (Little *et al.*, 2014) and a subset containing the 344 sequences that were obtained between 2005 and 2014. For both of these datasets, we inferred maximum-likelihood (ML) phylogenetic trees using the ModelFinder Plus feature (Kalyaanamoorthy *et al.*, 2017) of IQ-TREE (Chernomor *et al.*, 2016). We then removed outgroups from the tree inferred from the full 674 sequence dataset and used LSD (To *et al.*, 2016) to estimate the time of the most recent common ancestor (tMRCA) and the per-year mutation rate distribution.

The tMRCA was estimated at 1980. The mutation rate was estimated as 0.0012 with a standard deviation of roughly 0.0003, so to match these properties, we sampled mutation rates for each branch independently from a truncated Normal random variable from 0 to infinity with a location parameter of 0.0008 and a scale parameter of 0.0005 to scale branch lengths from years to expected number of per-site mutations.

In our simulations, a single viral lineage from each individual was sampled at the end time of the epidemic (10 years). The viral phylogeny in unit of time (years) was then sampled under a coalescent model with logistic viral population growth using the same approach as the the PANGEA-HIV methods comparison exercise, setting the initial population to 1, the per-year growth rate to 2.851904, and the time back from present at which the population is at half the carrying capacity (`v.T50`) to −2 (Ratmann *et al.*, 2017). Each seed individual is the root of an independent viral phylogenetic tree, and these trees were merged by simulating a seed tree with one leaf per seed node under a non-homogeneous Yule model (Le Gat, 2016) scaled such that its height equals 25 years to match the 1980 estimate using SD. The rate function of the non-homogeneous Yule model was set to 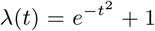 to emulate short branches close to the base of the tree (see comparison to other functions in Fig. S1).

##### Sequence data

We sampled a root sequence from a profile Hidden Markov Model (HMM) generated from the San Diego MSA using HMMER (Eddy, 1998). We evolved it down the scaled viral phylogenetic tree under the GTR+Γ model using Seq-Gen (Rambaut and Grass, 1997) with parameters inferred by IQ-TREE (Table S5).

##### Varying parameters

For San Diego, we explore four parameters (Table 1). For the contact network, in addition to the BA model, we used the ER (Bollobas, 1984) and WS (Watts and Strogatz, 1998) models. We also varied the expected degree (𝔼_*d*_) of individuals in the contact network between 2 and 16 (Table 1). For seed selection, we also used “Edge-Weighted,” where the probability that an individual is chosen is weighted by the individual’s degree. For each selection of contact network model, 𝔼_*d*_, and seed selection method, we study multiple rates of starting ART (expressed as 𝔼_*ART*_). In our discussions, we focus on 𝔼_*ART*_, a factor that the public health departments can try to impact. Increased effort in testing at-risk populations can decrease the diagnosis time, and the increased diagnosis rate coupled with high standards of care can lead to faster ART initiation. Behavioral intervention could in principle also impact degree distribution, another factor that we vary, but the extent of the effectiveness of behavioral interventions is unclear (Kelly *et al.*, 1991).

**Table 1:** Simulation parameters (base parameters in bold)

| Parameter | Parameter Values |
| --- | --- |
| Contact Network Model | **Barabási-Albert**, Erdős-Rényi, Watts-Strogatz |
| Expected Degree($\mathbb{E}_d$) | 2, **4**, 8, 16 |
| Seed Selection | **Random**, Edge-Weighted |
| Mean time to ART ($\mathbb{E}_{ART}$) | $\frac{1}{8}$, $\frac{1}{4}$, $\frac{1}{2}$, **1**, 2, 4, 8 (years) |

##### Uganda simulations

Our simulations with Uganda followed a similar approach to the base model used for San Diego but with different choices of parameters, motivated by Uganda. For inferring the reference phylogeny and mutation rates, we used a dataset of all 893 HIV-1 subtype D *pol* sequences in the Los Alamos National Laboratory (LANL) HIV Sequence Database that were sourced from Uganda and that were obtained between 2005 and 2014. All other Uganda parameters were motivated by McCreesh *et al.* (2017), and the following are key differences from the San Diego simulation. The contact network had 10,000 total individuals (a regional epidemic), and 1,500 individuals were randomly selected to be seeds. For epidemiological parameters, we assumed the expected time to begin as well as stop ART to be 1 year (McCreesh *et al.*, 2017). A comprehensive list of simulation parameters can be found in Tables S4 and S5.

#### 2.2.2 Transmission network reconstruction methods

We compare two HIV network inference tools: HIV-TRACE (Kosakovsky Pond *et al.*, 2018) and TreeCluster (Moshiri, 2018). HIV-TRACE is a widely-used method (Rose *et al.*, 2017; Wertheim *et al.*, 2017; Pérez-Losada *et al.*, 2017) that clusters individuals such that, for all pairs of individuals *υ* and *ν*, if the Tamura and Nei (1993) (TN93) distance is below the threshold (default 1.5%), *υ* and *ν* are connected by an edge; each connected component forms a cluster. When we ran HIV-TRACE, we skipped its alignment step because we did not simulate indels. TreeCluster clusters the leaves of a given tree such that the pairwise path length between any two leaves in the same cluster is below the threshold (default 4.5%), the members of a cluster define a full clade, and the number of clusters is minimized. Trees given to TreeCluster were inferred using FastTree 2 (Price *et al.*, 2010) under the GTR+Γ model. We used FastTree 2 because using IQ-TREE on these very large datasets (up to 80,000 leaves) was not feasible. TreeCluster is similar in idea to Cluster Picker (Ragonnet-Cronin *et al.*, 2013), which uses sequence distances instead of tree distances (but also considers branch support). We study TreeCluster instead of Cluster Picker because of its improved speed. Our attempts to run PhyloPart (Prosperi *et al.*, 2011) were unsuccessful due to running time.

#### 2.2.3 Measuring the predictive power of clustering methods

We now have two sets of clusters at the end of the simulation process (year 10): one produced by HIV-TRACE and one by TreeCluster. Let *C*^*t*^ denote the clustering resulting from removing all individuals infected after year *t* from a given final clustering *C*^10^, let 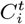 denote a single *i*-th cluster in clustering *C*^*t*^, and let 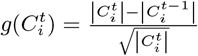 denote the growth rate of a given cluster 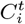 (Wertheim *et al.*, 2018). We then compute the average number of individuals who were infected between years 9 and 10 by the “top” 1,000 individuals (roughly 5% of the total infected population) who were infected at year 9, where we choose top individuals by sorting the clusters in *C*^9^ in descending order of 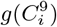 (breaking ties randomly) and choosing 1,000 individuals in this sorting, breaking ties in a given cluster randomly if needed (e.g. for the last cluster needed to reach 1,000 individuals). As a baseline, we compute the average number of individuals who were infected between years 9 and 10 by *all* individuals, which is equivalent (in expectation) to a random selection of 1,000 individuals. Our metric, therefore, measures the risk of transmission from the selected 1,000 individuals. Our motivation for this metric is to capture whether monitoring cluster growth can help public health intervention efforts with limited resources in finding individuals with a higher risk of transmitting.

## 3 Results

### 3.1 Comparison to real phylogenies

To compare data simulated by FAVITES to real data, we use the aforementioned San Diego and Uganda phylogenies. Since the trees on real data are inferred trees (as opposed to true trees), we compare them to inferred trees on simulated data (built using FastTree 2 as running IQ-TREE on simulated data was not feasible). We randomly subsample the simulated dataset to match the number of sequences in the corresponding real dataset (344 for San Diego; 893 for Uganda).

For San Diego, the mean patristic distance between sequences on inferred trees is respectively 0.087 and 0.089 for the real and base simulated datasets. The distributions of pairwise distances among inferred trees of real and simulated datasets have similar shapes, but distances from real data have higher variance (Fig. 3a). To quantify the divergence between the real and simulated distributions, we use the Jensen-Shannon Divergence (JSD), a number between 0 and 1 with 0 indicating a perfect match (Lin, 1991). The JSD is only 0.023 for trees inferred from the San Diego base parameters (Table S6). The Uganda simulations have a larger divergence (Fig. 3a) between real and simulated distributions (JSD: 0.082), with simulated data showing higher mean distances (means: 0.075 and 0.097). We observe similar patterns when we compute pairwise distances directly from sequences and apply phylogenetic correction using the JC69+Γ model (Table S6; Fig. S4). For all simulated datasets, the true trees have lower variance in pairwise distances compared to estimated trees; this is consistent with the stochasticity of sequence evolution and the added variance due to the inference uncertanity.

**Figure 3:**
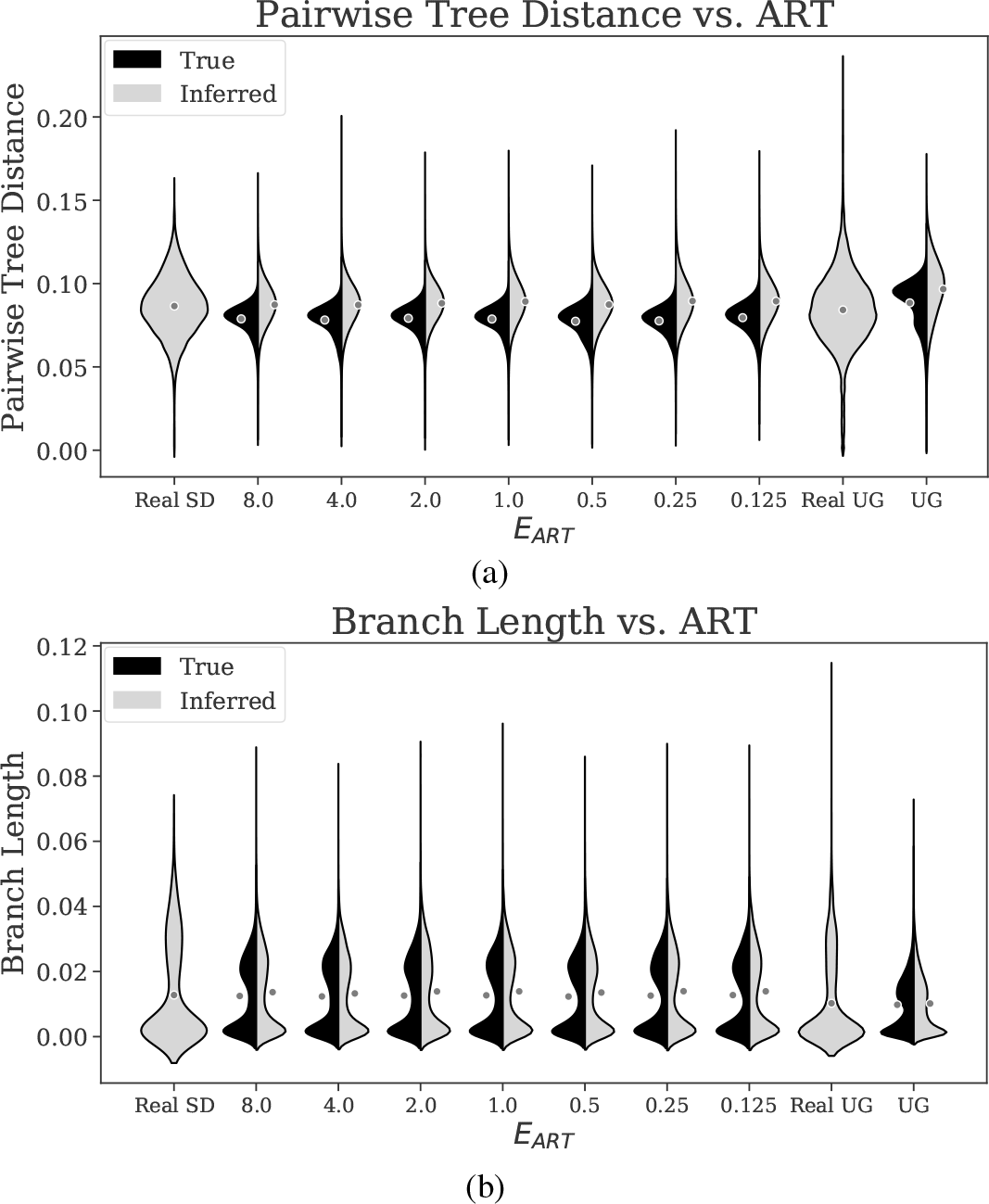
Kernel density estimates of the distributions of (a) patristic distances (path length) between all pairs of sequences and (b) branch lengths of real and simulated datasets for the San Diego (SD) and Uganda (UG) datasets. Averages are shown as dots (Fig. S3): Black denotes distributions computed from true (simulated) trees and grey denotes distributions computed from trees inferred from sequences (IQ-TREE for real and FastTree 2 for simulated data). Note that real data only have inferred pairwise distances and branch lengths, as true branch lengths are not known. 𝔼_*ART*_ is the expected time to start ART.

Our simulated trees, like real trees, include clusters of long terminal branches and short internal branches, especially close to the root (Fig. S2). The branch length distributions are bimodal, with one peak close to 0 and another between 0.01 and 0.03 (Fig. 3b). However, the second mode for the real trees is larger than the second mode of real data; for example, for San Diego, the second peak is at 0.030 for real data and 0.023 for base simulated data. The JSD divergence between branch length distributions of real and simulated trees are 0.102 for San Diego (base) and 0.119 for Uganda. The distribution of branch lengths on true trees (as opposed to inferred trees) has a similar shape (Fig. 3b) but a shorter tail of long branches and a reduced JSD compared to real data (e.g., 0.044 for base San Diego; see Table S6).

#### Sensitivity to parameters

Even though mean branch lengths can change (between 0.0053 and 0.0080) as a result of changing 𝔼_*ART*_ and 𝔼_*d*_ (Fig. S3), the overall distributions remain quite stable (Figs. 3b and S4). Similarly, patristic distances are not sensitive to E_ART_ (Fig. 3ab) nor to 𝔼_*d*_ (Fig. S4). In terms of branch lengths, the divergence from the real data changes only marginally as 𝔼_*d*_ and 𝔼_*ART*_ change (Table S7). While the distributions are stable with respect to these epidemiological parameters, they are sensitive to others. For example, results are sensitive to the model of mutation rates. We draw mutation rates from a Truncated Normal distribution (fitted to real data) and obtain close matches to real data. However, other distributions (e.g. Exponential) yield significant deviation from real distributions (Fig. S4). Because of these deviations, we have only used the truncated normal distributions for mutation rates everywhere.

### 3.2 Impact of parameter choices on the epidemiology

#### Infected population

The number of infected individuals increases with time and the rate of growth is faster for larger 𝔼_*ART*_ values (Fig. S5). For all tested values of 𝔼_ART_, the number of infected individuals grows close to linearly (Pearson *r* ≥ 0.966), indicating that the large at-risk population has not saturated in the 10-year simulation period. As 𝔼_*ART*_ decreases from 8 years to 1/8 years, the total number of infected individuals at the end of the simulation keeps decreasing (Fig. 4a). For example, with degree 4, the average final number of infected individuals in the 10 year period is 6686, 4134, and 1273 with 𝔼_*AR*T_ set to 1, 1/2, 1/8 year, respectively.

The model of contact network and the model of choosing the seed individuals have only marginal effects on these outcomes. Edge-weighting the seed selections yields a slightly higher (at most 12%) total number of infected individuals than the random selection (Table S7). The BA model of contact network leads to a slightly higher infection count when compared to the ER (at most 7%) and WS (at most 8%) models (Figs. S6), but these differences are marginal compared to impacts of 𝔼_*ART*_ and 𝔼_*d*_ (which, when changed, leads to 43% and 152% change, respectively, in the number of infected people compared to the base parameters). Finally, Uganda simulations lead to higher infection count (64% versus 45%) compared to San Diego (Table S7).

**Figure 4:**
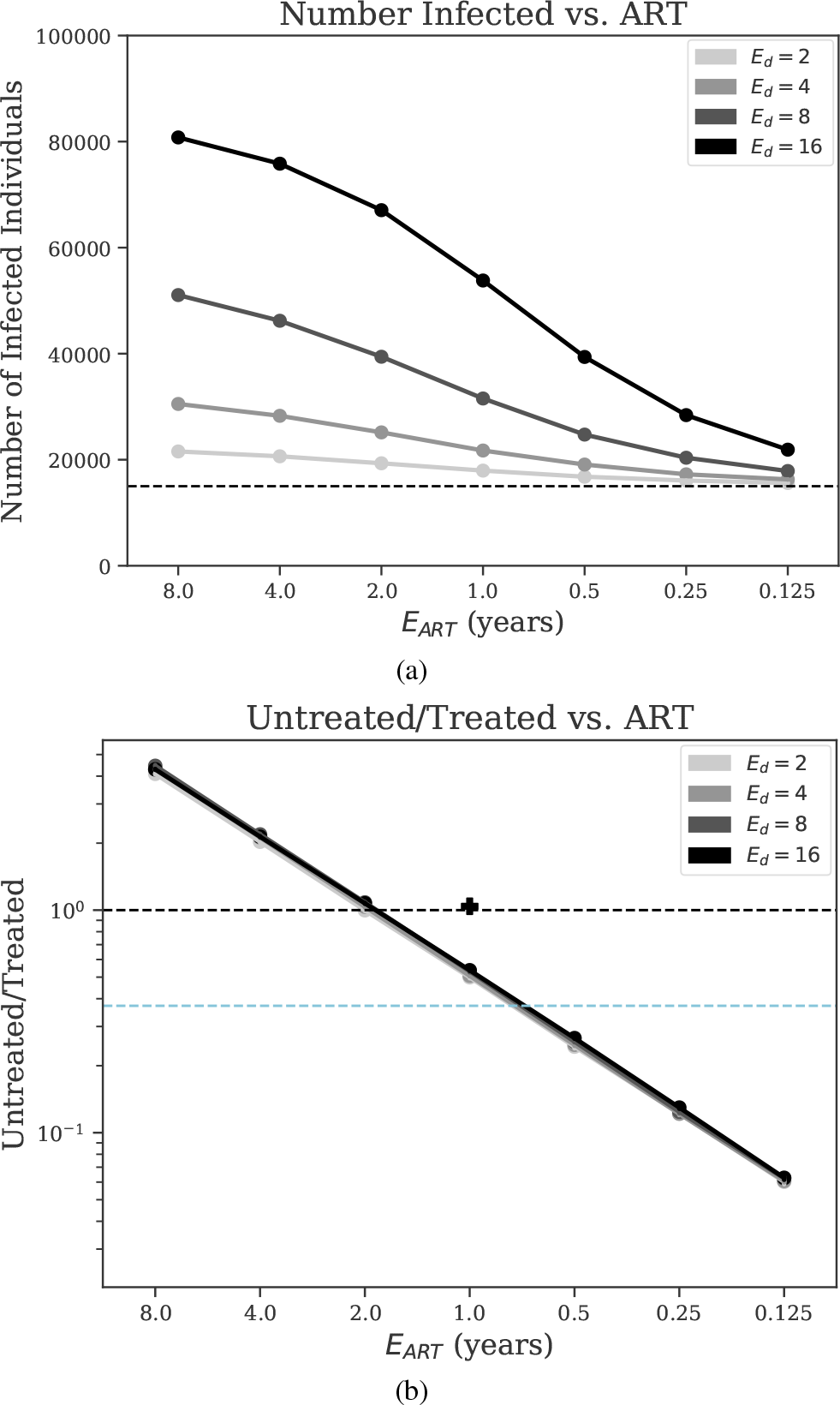
Sensitivity analysis of epidemiological outcomes. We show (a) the total number of infected individuals, and (b) the ratio of the number of untreated vs. the number of treated individuals (log-scale), vs. expected time to begin Antiretroviral Therapy (𝔼_*ART*_) for the Barabási-Albert model with various mean contact numbers (𝔼_*d*_) with all other parameters set to base values. Untreated/treated = 1 is shown as a dashed black line, and the value of untreated/treated corresponding to the “90-90-90” goal (UNAIDS, 2014) is shown as a dashed blue line ((1 − 0.9^3^)/0.9^3^ ≈ 0.37). The Untreated/Treated value corresponding to the simulated Uganda dataset has been shown as a + symbol on (b).

#### Treated population

The ratio of untreated to treated individuals is a function of 𝔼_*ART*_ but not 𝔼_*d*_ (Fig. 4b). Note that this ratio remains constant (at most 14.7% change) after year 4, has small changes in year 1 to 4, and experiences an initial period of instability for about 1 year (Fig. S5), likely because all seeds are initially AU. With 𝔼_*ART*_ = 1 years, the ratio is on average 0.507 after year 2; decreasing/increasing 𝔼_*ART*_ reduces/increases the portion of untreated people. The 90-90-90 campaign by UNAIDS (2014) aims to have 90% of the HIV population diagnosed, of which 90% should receive treatment, of which 90% (i.e., 72.9% of total) should be virally suppressed. Reaching the 90-90-90 goals in the epidemic we model here requires 𝔼_*ART*_ between 1/2 and 1 year (assuming that lack of viral suppression is fully attributed to lack of adherence). These results are stable with respect to model of contact network, 𝔼_*d*_, and seed selection approach (Figs. 4b and S7). The only model choice that had a noticeable effect on the results is the use of the ER network model, which led to an increase in Untreated/Treated for 𝔼_*d*_≤ 4 (Fig. S7). We note that our simulated Uganda epidemic had twice the ratio of Untreated/Treated compared to base San Diego (Table S7).

### 3.3 Evaluating inference methods

#### Phylogenetic error

From simulated sequences, we inferred trees under the GTR+Γ model using FastTree 2 (Price *et al.*, 2010), and we computed the normalized Robinson-Foulds (RF) distance (i.e., the proportion of branches included in one tree but not the other (Robinson and Foulds, 1981)) between the true trees and their respective inferred trees (Fig. S8). For all model conditions, the RF distance is quite high (0.36-0.58 for San Diego and 0.25-0.40 for Uganda). However, we note that our datasets include many extremely short branches, defined here as those where the expected number of mutations along the branch across the entire sequence length is lower than 1. In our simulations, we have between 16% and 30% of branches that are extremely short (Fig. S8) and therefore hard to infer.

#### Clustering methods

We measure the number of new infections caused by each person in the clusters with the highest growth rate and compare it with the same value for the total population (Fig. 5). Over the entire population, the average number of new infections caused by each person between years 9 and 10 is 0.028 for our base parameter settings. The top 1,000 people from the fastest growing TreeCluster clusters, in contrast, infect on average 0.066 new people. Thus, the top 1000 people chosen among the growing clusters according to TreeCluster are more than twice as infectious as a random selection of 1000 individuals. HIV-TRACE performs even better than TreeCluster, increasing the per capita new infections among top 1,000 individuals to 0.097 for base parameters, a 3.46× improvement compared to the population average. As 𝔼_*ART*_ decreases, the total number of per capita new infections reduces; as a result, the positive impact of using clustering methods to find the growing clusters gradually diminishes (Fig. 5). Conversely, reducing 𝔼_*ART*_ leads to further improvements obtained using TreeCluster versus random selection and using HIV-TRACE versus TreeCluster.

**Figure 5:**
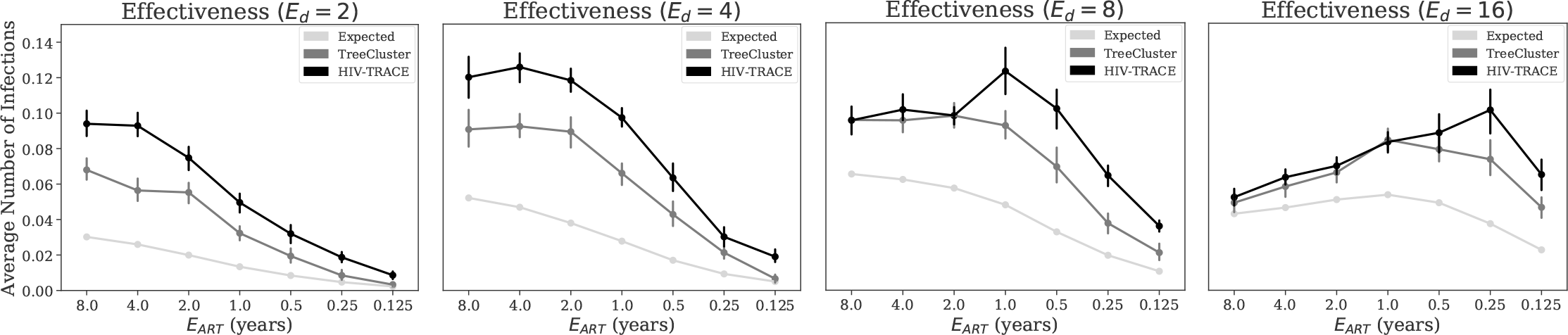
The effectiveness of clustering methods in finding; high risk individuals. The average number of new infections between years 9 and 10 of the simulation caused by individuals infected at year 9) in growing clusters. We select 1,000 individuals from clusters, inferred by either HIV-TRACE or TreeCluster, that have the highest growth rate (ties broken randomly). As a baseline control, the average number of infections over all individuals (similar’ to expectations under a random selection) is shown as well. For a cluster with *n*_*t*_ members at year *t*, growth rate is defined as 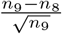. The columns show varying expected degree (i.e., number of sexual partners), and all other parameters are their base values.

Changing 𝔼_*d*_ also impacts the results (Fig. 5). When 𝔼_*d*_ = 2, slowing the epidemic down compared to the base case, both methods remain better than random, and HIV-TRACE continues to outperform TreeCluster. However, when 𝔼_*d*_ is increased, the two methods first tie at 𝔼_*d*_ = 8, and at 𝔼_*d*_ = 16, TreeCluster becomes slightly better than HIV-TRACE for most E_art_ values (Fig. 5). The advantage compared to a random selection of individuals is diminished (improvements never exceed 70%) when the epidemic is made very fast growing by setting 𝔼_*ART*_≥ 2 and 𝔼_*d*_ = 16.

## 4 Discussion

Our results demonstrated that FAVITES can simulate under different models and can produce realistic data. A comparison of the fit between real and simulated data for Uganda and San Diego points to the importance of data availability. For San Diego, where more studies have been done and more sequence data were available, the fit between simulated and true data was generally good (Table S6). For Uganda, we had to rely on several sources (e.g., data from McCreesh *et al.* (2017) and LANL), and we had a reduced fit between simulations and real data. Increased gathering and sharing of data, including sequence data, can in future improve our ability to parameterize simulations.

Although we only explored viral epidemics, FAVITES can easily expand to epidemics caused by other pathogens for which molecular epidemiology is of interest (Azarian *et al.*, 2014). We also showed that TreeCluster and HIV-TRACE, when paired with temporal monitoring, can successfully identify individuals most likely to transmit, and HIV-TRACE performs better than TreeCluster under most tested conditions. The ability to find people with increased risk of onward transmission is especially important because it can potentially help public health officials better spend their limited budgets for targeted prevention (e.g. pre-exposure prophylaxis, PrEP) or treatment (e.g. efforts to increase ART adherence).

We studied several models for various steps of our simulations, but we did not exhaustively test all models: FAVITES currently includes 21 modules and a total of 169 implementations (i.e., specific models) across them, and testing all model combinations is infeasible. To simulate San Diego and Uganda, we aimed to choose the most appropriate set of 21 sub-models available in FAVITES to create the end-to-end simulations. Each of these 21 sub-models has its own limitations, as models inevitably do. However, it must be noted that limitations resulting from model assumptions are limitations of the specific example simulation experiment we performed in this manuscript, rather than limitations of the framework: FAVITES is designed specifically to be flexible, allowing the use of different models for different steps. If better models are developed for each of these 21 modules, they can be easily incorporated. Like all statistical modeling, appropriate choice of model assumptions is essential to the interpretation of the simulation results, and it is important for the user to choose models appropriate to their specific epidemic of interest. To aid users, our extensive documentation provides descriptions for each module implementation and we provide model validation scripts.

For the simulation of HIV epidemics, novel statistical models can be created to address the unrealistic assumptions. For example, our contact network remains unchanged with time, whereas real sexual networks are dynamic. Our transmission model does not directly model effective prevention measures such as PrEP. Our sequences include substitutions, but no recombination. Moreover, the models of sequence evolution we used ignore many evolutionary constraints across sites. We also ignored infections from outside the network; (viral migration), assumed full patient sampling, and we sampled all patients at; the end time as opposed to varied-time sampling. While these and other choices may impact results, we note that our goal here was mainly to show the utility of FAVITES. We leave an extensive study of the impact of each of these factors on the results to future studies. Importantly, new models with improved realism to address these issues can easily be incorporated, and continued model improvement is a reason why we believe flexible frameworks like FAVITES are needed.

We observed relatively high levels of error in inferred phylogenies. This is not surprising given the low rate of evolution and length of the *pol* region (which we emulate). Further, our phylogenies include many super-short branches, perhaps due to our complete sampling;. Many transmission cluster inference tools (e.g. PhyloPart, Cluster Picker, and TreeCluster) use phylogenies during the inference process and thus may be sensitive to tree inference error. Other tools like HIV-TRACE do not; attempt to infer a full phylogeny (only distances). The high levels of tree inference error may be partially responsible for the relatively grower performance 0f. TreeCluster compared to HIV-TRACE. Nevertheless, TreeCluster had higher per capita new infections in its fastest growing clusters than the population average, indicating that the trees, although imperfect, still include useful signal about the underlying transmission histories).

Using FAVITES, we compared TreeCluster and HIV-TRACE in terms of their predictive power, and our results complement studies on real data (Rose *et al.*, 2017). Nevertheless, our simulations study has some limitations that should be kept in mind. A major limitation is that both methods we tested use a distance threshold internally for defining clusters. The specific choice oi threshold defines a trade-off between cluster sensitivity and specificity, and the trade-off will impact cluster compositions. T he best choice of the threshold is likely a function of epidemiological factors, and the default thresholds are perhaps optimal for certain epidemiological conditions, but; not; others. For example, we observed that, for a minority of our epidemiological settings, TreeCluster is more effective than HIV-TRACE in predicting growing clusters. A thorough exploration of all epidemiological parameters and method thresholds is left for future studies. On a practical note, FAVITES can enable public health officials to simulate conditions similar to their own epidemic and pick the best method/threshold tailored to their situation.

The approach we used for evaluating clustering methods, despite its natural appeal, is not the only possible measure. For example, the best way to choose high-risk individuals given clustering results at one time point or a series of time points is unclear. We used a strict ordering based on square-root-normalized cluster growth and arbitrary tie-breaking, but many other metrics and strategies can be imagined (Wertheim *et al.*, 2018). For example, we may want to order individuals within a cluster by some criteria as well and choose certain number of people per cluster inversely proportional to the growth rate of the cluster. We simply chose 1,000 people to simulate a limited budget, but perhaps reducing/increasing this threshold gives interesting results. A thorough exploration of the best method for each budget is beyond the scope of this work. Similarly, we leave a comprehensive study of the best strategies to allocate budgets based on the results of clustering and better ways of measuring effectiveness, to future work.

